# Stochastic Boolean Model of Death Signaling in MCF7:5C Predicts Cell Death Inducers and Inhibitors

**DOI:** 10.64898/2026.09.05.749284

**Authors:** Kittisak Taoma, Teeraphan Laomettachit, Surojeet Sengupta, Yue Wang, Robert Clarke, Pavel Kraikivski

## Abstract

Disrupted cell death signaling contributes significantly to the inappropriate survival of cancer cells, enabling them to evade the apoptotic processes that normally eliminate damaged or abnormal cells. To assess the impact of disrupted cell death signaling on cancer cell survival, we developed a continuous-time stochastic Boolean model of apoptosis signaling in MCF7:5C human breast cancer cells that are resistant to estrogen deprivation and undergo apoptosis in response to estrogen. The model was calibrated to replicate the dynamics of key apoptosis regulators in MCF7:5C cells before and after 17β-estradiol (E2) treatment. Subsequently, the calibrated model was used to predict both the single and double perturbations that inhibit cell death in estradiol-treated MCF7:5C cells, and additional single interventions capable of reinducing apoptosis. For example, strains with single-gene deletion of PERK, eIF2α, CHOP, or ATF4 enabled E2-treated MCF7:5C cells to evade apoptosis. However, additional inhibition of SRC, PI3K, ESR1, CEBPB, or MAPK8 restored apoptotic responses in these resistant cells. We identified 30 resistant cell strains with double mutations and 506 drug-target combinations capable of inducing four distinct modes of cell death: direct CASP7 activation, extrinsic apoptosis, intrinsic apoptosis, and combined intrinsic/extrinsic apoptosis. Overall, our model successfully explains the drug response behavior of MCF7:5C cells, predicts mechanisms of drug resistance, and suggests treatment strategies to overcome this resistance.

## Introduction

Breast cancer is a heterogeneous disease with several subtypes, each with a distinct molecular profile. The luminal subtype, also often referred to as hormone receptor-positive (HR+) accounts for ∼70% of all breast tumors and is characterized by the presence of progesterone (PR) and/or estrogen (ER) receptors ^1^. A common strategy for treating HR-positive breast cancer involves endocrine therapy ^2^. The treatment protocols include three classes of drugs: (i) selective estrogen receptor modulators (SERMs), such as tamoxifen ^3^, (ii) selective estrogen receptor downregulators (SERDs), such as fulvestrant ^4^, and (iii) aromatase inhibitors, such as anastrozole and letrozole ^5^.

Despite the significant success of endocrine or cytotoxic therapies ^6,7^, drug resistance remains a major obstacle to curing cancer patients ^8^. Therapeutic interventions involving drug combinations have been proposed to mitigate the development of drug resistance ^9^. Combination therapies can offer enhanced efficacy, allow for dose reduction, decrease toxicity, and improve long term survival for patients ^10^. However, identifying drug targets for combination therapies can be both costly and time-consuming, involving everything from target discovery to clinical trials ^11^. To make this process more efficient, mathematical models that account for factors contributing to drug resistance can be applied. These models can test various treatment strategies and predict the most effective protocols to enhance drug efficacy and improve patient survival.

As mathematical models have long been used to infer regulatory networks ^12^, they have been successfully used to identify potential resistance pathways and therapeutic targets in both HR+ ^13–15^ and triple negative breast cancer (lacks ER, PR and HER2) ^15–17^. However, it remains uncertain whether these approaches can effectively uncover potential drug resistance mechanisms and optimal drug combinations in endocrine-resistant breast cancer cell lines. For example, it is counterintuitive that E2 treatment can reduce breast cancer incidence in some postmenopausal breast cancers resistant to aromatase inhibitors, as observed in clinical trials ^18^. A similar response to E2 treatment is observed in the MCF7:5C cell line. MCF-7:5C cells are an estrogen receptor-positive (ER+) variant of the parental MCF-7 breast cancer cell line that was developed through long-term estrogen deprivation and exhibits estrogen-independent growth and resistance to antiestrogen treatment. Thus, MCF-7:5C cells provide an established model of endocrine resistance and have been used to investigate mechanisms associated with resistance to estrogen-deprivation therapies, including aromatase inhibitors. Notably, in contrast to most ER+ breast cancer cell lines, including parental MCF-7 cells, E2 treatment of MCF-7:5C cells induces growth inhibition and apoptosis ^19,20^. This distinctive phenotype provides a useful model for studying estrogen-induced apoptosis in endocrine-resistant breast cancer and for identifying druggable molecular targets ^21^ and therapeutic combination strategies that may help overcome endocrine resistance and limit tumor growth ^22,23^.

While many treatment protocols can effectively shrink tumors, drug resistance often develops due to the emergence of new resistant subclones under therapeutic pressure, either by adaptation (acquired resistance) or from pre-existing resistant populations ^24^. The molecular mechanisms driving acquired resistance to anticancer therapies often involve the activation of pro-survival signaling networks and anti-apoptotic pathways ^25^. Therefore, understanding how alterations in cell death regulatory mechanisms diminish treatment efficacy is essential for identifying complementary therapeutic targets capable of reducing resistance. Developing a mathematical model that simulates the balance between pro-survival and pro-death signaling in cancer cells could facilitate the systematic evaluation of potential resistance mechanisms and support the design of treatment strategies aimed at preventing, delaying, or overcoming therapeutic resistance.

In this work, we propose a mathematical model to study how to effectively drive endocrine-resistant cancer cells toward irreversible cell death. We calibrate the model using experimental data from MCF7:5C cancer cells to replicate resistance to E2-induced cell death in breast cancer patients. Since the apoptosis pathway is critical for E2-induced cell death in MCF7-5C cells ^26^, our model focuses on the molecular mechanisms of E2-induced apoptosis (see Fig 1). The model is used to identify perturbations that inhibit cell death in the presence of E2 treatment. These perturbations simulate molecular modifications, including acquired gene mutations or epigenetic changes, which alter gene expression. Cancer cells with modifications that prevent E2-induced cell death are classified as having an E2-resistant phenotype. We then use the model to discover new molecular targets that can induce cell death in these resistant cells. Our findings can help develop new therapeutic strategies and drug combinations to effectively treat E2-resistant cancer.

**Fig 1.**
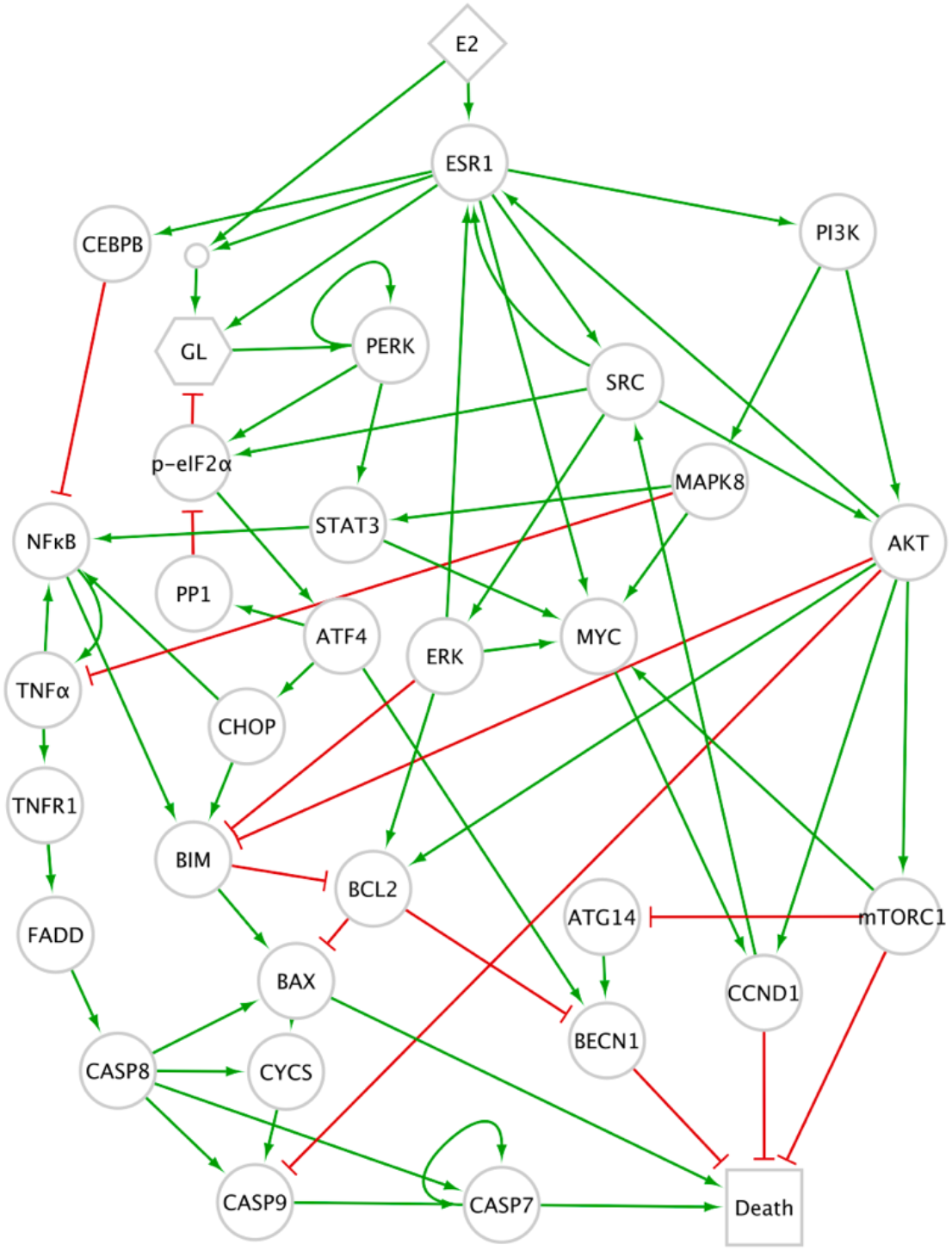
Influence diagram representing the modeled regulation of E2-induced apoptosis in MCF7:5C cells. The diagram illustrates a regulatory network with nodes connected by edges that represent inhibition (red lines with blunt ends), activation (green lines with barbed ends), and “AND” gate (small circle). Diamond-shaped nodes represent ligands, circles represent proteins, squares represent progress variables, and rectangles represent readout variables. The prefix “p-” denotes the phosphorylated state of a protein.

## Methods

### Network reconstruction of E2-induced apoptosis in MCF7:5C cells

The molecular signaling network regulating E2-induced apoptosis in MCF7:5C cells was reconstructed through a multi-step curation process. First, a core network was assembled based on available experimental data from MCF7:5C cells. This initial network captured the dual role of the PERK-mediated stress pathway, which functions as a molecular switch governing the transition of MCF7:5C cells from growth to apoptosis in response to E2 treatment ^27^. Next, the core network was expanded and refined by incorporating additional nodes and regulatory interactions from the SIGNOR 4.0 database ^28^. To ensure biological relevance to the MCF7:5C cell line, only those interactions that improved the model’s ability to reproduce experimentally observed MCF7:5C responses were retained. Expansion and refinement were conducted iteratively through a trial-and-error approach using model calibration and validation against available experimental data. This iterative process continued until the final network was able to accurately reproduce and explain the observed E2-induced apoptotic responses in MCF7:5C cells. The final regulatory network describing E2-induced apoptosis is presented in Fig 1. A complete list of network nodes and interactions, together with the references used for their curation, is provided in S1Table.

### Boolean modeling framework

The regulatory network shown in Fig 1 was modeled using a time-continuous stochastic Boolean framework ^29^, which enables systematic analysis of the dynamic behavior of all network components. The state of each node *i* in the regulatory network at a given time *t* is represented by a Boolean variable *X*_*i,t*_. The potential change in *X*_*i,t*_ depends on the states of nodes 1 through *n*, described by the corresponding variables *X*_1,*t*_, *X*_2,*t*_, …, *X*_*n,t*_, which either activate or inhibit node *i*, as shown in Fig 1. Our model captures the dynamics of 29 proteins; therefore, *n*=29. The influence of nodes 1 through *n* on node *i* is represented by the following weighted sum:

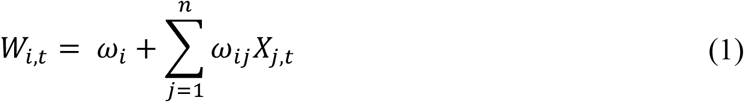

where *ω*_*i*_ and *ω*_*ij*_ are adjustable coefficients determining the threshold at which the state of the node *i* changes. Coefficients were chosen from values ranging from −4 to 4 at 0.5 increments (see S2 Table) to fit the experimental data and reproduce the observed sequence of protein activation and inactivation events, as described in the model calibration subsection below. All *W*_*i*_ functions used in our Boolean model are summarized in Table 1. Inhibitory regulatory interactions are represented by negative signs directly within the update equations; therefore, all parameter values *ω*_*i*_ and *ω*_*ij*_ presented in **Table S2** are positive.

**Table 1.**
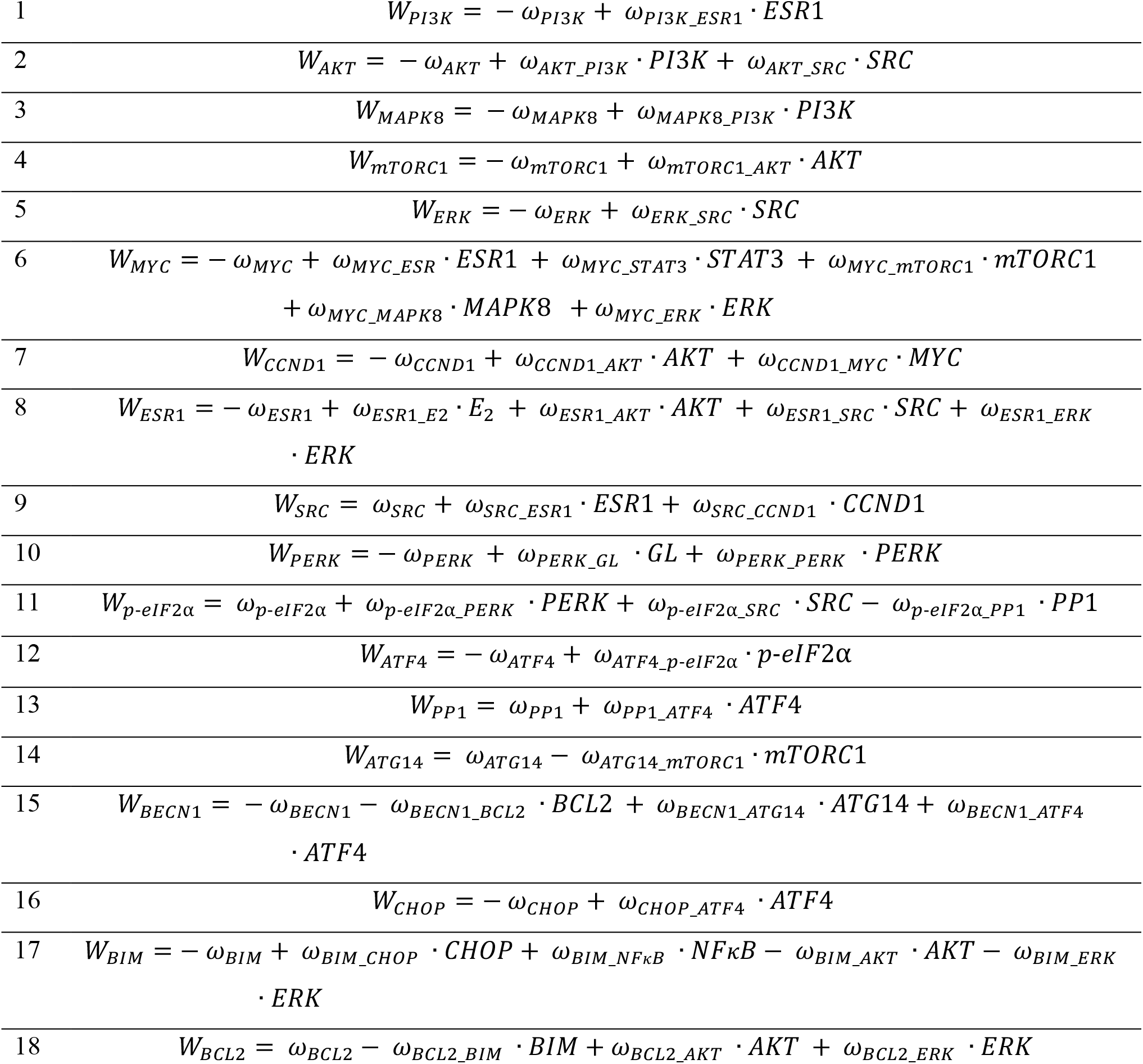
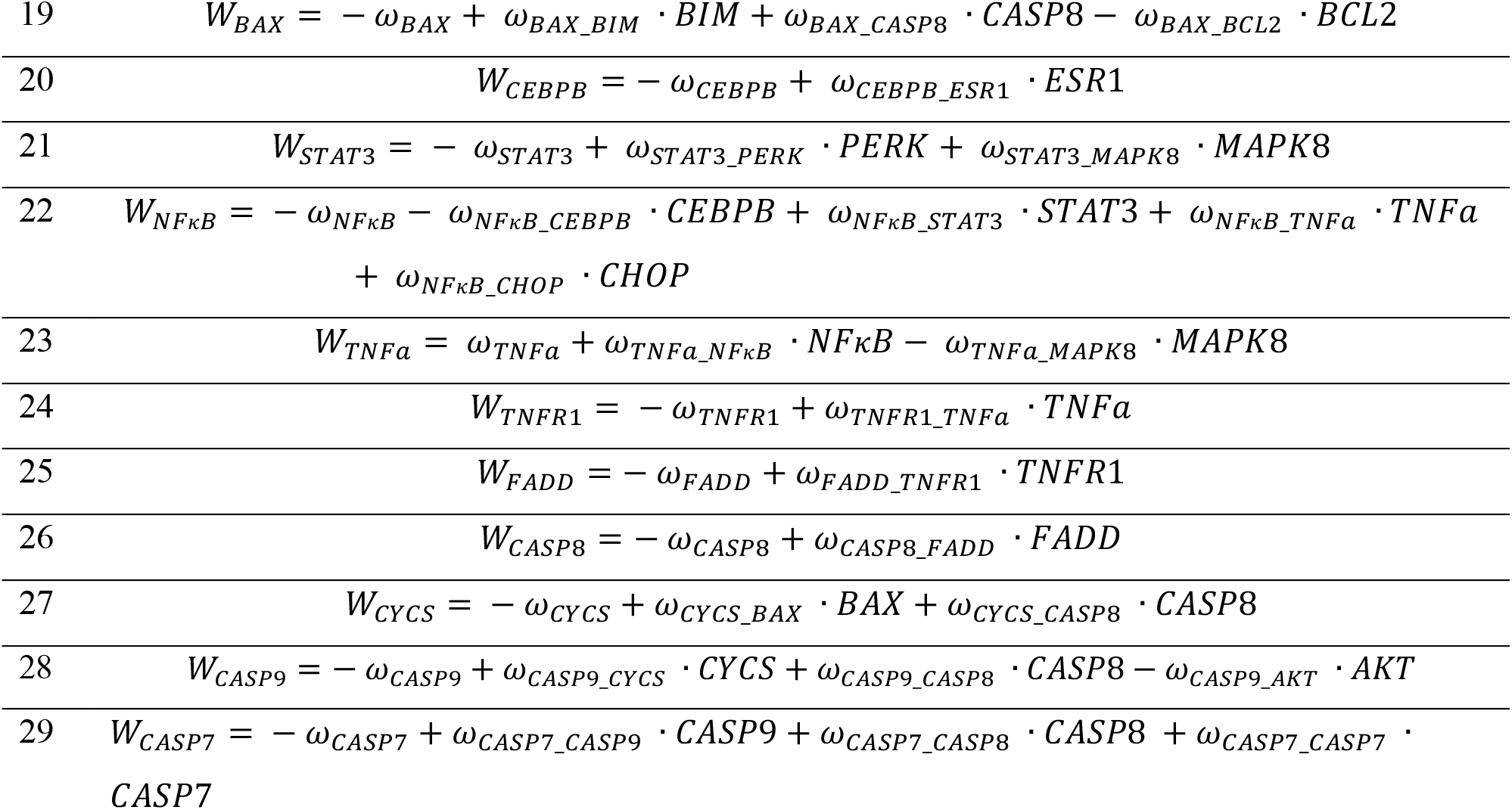
Update functions (*W*_*i*_) for the 29 proteins in the Boolean network of E2-induced apoptosis in MCF7:5C cells.

The functions *W*_*i,t*_ are binarized using the Heaviside function to determine the potential change of all variables 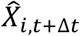 at time step *t* + Δ*t* as follows:

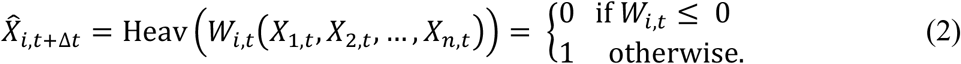

The hat above the variable indicates the potential change, implying that the new value may or may not be set, depending on whether the variable is selected for updating. An asynchronous update scheme was used, meaning that only one variable, 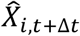, is updated at each time step *t* + Δ*t*. The selection of which variable to update is based on the following propensity functions:

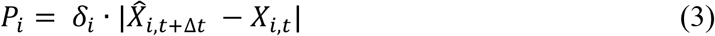

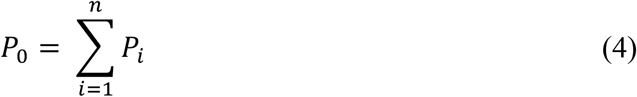

where *δ*_*i*_ is a constant representing the probability per unit time that *X*_*i*_ undergoes a potential change. The propensity (*P*_*i*_) is proportional to the magnitude of the predicted change in (*X*_*i*_), while (*P*_0_) denotes the total propensity across all variables. In this work, we set all *δ*_*i*_ to 1 to ensure equal probability for updating all variables. The index of the selected variable to update at the next time step is determined as the smallest integer *j* that satisfies: 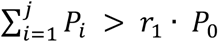, where *r*_1_ is a random number drawn from a uniform distribution between 0 and 1.

The time interval Δ*t* at which the variable change occurs is computed as:

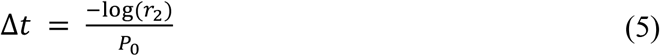

where *r*_2_ is another random number generated from a uniform distribution between 0 and 1, independently of *r*_1_.

To model the activation of apoptosis in MCF7:5C cells following prolonged exposure to E2, we introduced a time delay for the activation of CHOP and CASP8 proteins using a gamma distribution with the following probability density function:

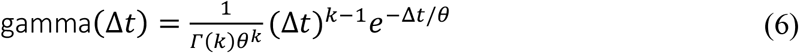

where the mean value is given by *κ* · *θ* and coefficient of variation is 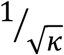. To align with experimental data on apoptosis activation, we set *κ* = 30 and *θ* = 1 for CHOP (mean = 30 min and CV = 0.18), and *κ* = 10 and *θ* = 1 for CASP8 (mean = 10 min and CV = 0.31).

To represent the amount of global protein synthesis, we introduced a progress variable, global protein loading (*GPL*). The increase in global protein synthesis was modeled using an exponential function *GPL*_*t*+Δ*t*_ = *GPL*_*t*_ · *e*^*λ*·Δ*t*^, where the protein synthesis rate (*λ*) is calculated as: 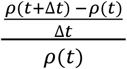. Here, *ρ*(*t*) and *ρ*(*t* + Δ*t*) are the time-dependent effects of E2-induced global protein synthesis in MCF7:5C cells, as detected by Western blot ^21^. Band intensities were quantified by digitizing the Western blot images using ImageJ software (Version 1.54m).

Decreases in protein synthesis induced by phosphorylation of the eukaryotic translation initiation factor eIF2*a* were modeled as: *GPL*_*t*+Δ*t*_ = *β* · *GPL*_*t*_, where the *β* denotes the protein synthesis reduction factor. The value of *β* was estimated from experimental measurements of the decrease in protein synthesis activity observed between 48 and 72 hours. The derivations of the parameters *λ* and *β* are explained in the S1 Text. Finally, the *GPL* was set to zero when the apoptosis executor (CASP7) is activated. In summary, changes in the global protein synthesis variable are governed by the following rules:

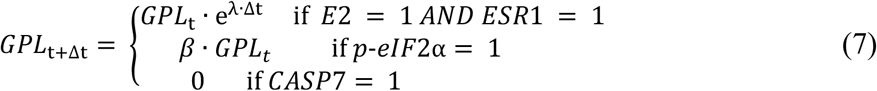

We convert the progress variable *GPL* to a Boolean variable *GL* by using Heaviside function:

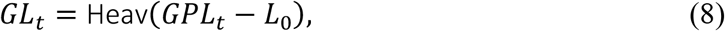

where *L*_0_ is a threshold used to determine the timing for PERK protein activation, which occurs along with an increase in the global protein synthesis.

We also used readout variables to monitor pro-death and pro-survival events in MCF7:5C cells. The activation of *CCND1* is used to detect G1/S transition, *mTORC1* activation monitors protein synthesis, *BECN1* is used to detect autophagy activation, and *CASP7* tracks the execution of apoptosis. To evaluate a “*Death*” score, we calculated the difference between pro-death readout variable (*CASP7*) and variables associated with cell proliferation, growth and survival (*CCND1, mTORC1, BECN1*). This difference was then passed through a Sigmoid function, as follows

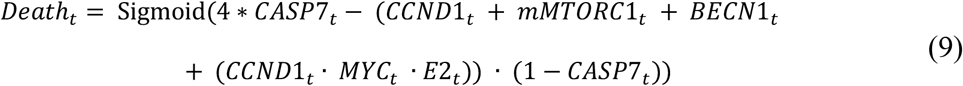

According to Equation (9), the death score approaches 1 when CASP7 protein is activated. In this case, pro-survival proteins that promote replication and growth do not contribute to the score. The *CCND*1_*t*_ · *MYC*_*t*_ · *E*2_*t*_ complex was included to capture the transient increase in cell proliferation induced by E2 prior to apoptotic cell death ^30^. However, when CASP7 is inactive (i.e., *CASP7 =* 0), the score is determined by the activity of the pro-survival proteins. Coefficient 4 in front of the CASP7 term is introduced to balance the combined contribution of the pro-survival terms, which collectively sum to 4 when all pro-survival proteins are fully active. The sigmoid function used in Equation (9) is defined as: 1/(1 + *e*^−2*x*^). This function approaches 0 when CASP7 is inactive and the pro-survival terms are fully active, and it approaches 1 as CASP7 activity increases, thereby capturing the switch-like behavior between survival and apoptotic signaling.

### Model calibration

The equations in Table 1 involve 90 coefficients, *ω*_*ij*_ and *ω*_*i*_, which were manually adjusted to align protein activation and inactivation events with experimental observations in both control and drug-exposed MCF7:5C cells. All data used for model calibration were obtained from the literature and are summarized in Tables 2 and 3. The values of all coefficients, *ω*_*ij*_ and *ω*_*i*_, for the calibrated model are provided in S2 Table. Simulated protein activities were recorded after 300 simulation time steps, at which point all protein activities had reached steady-state values. The simulation results from both unperturbed and drug-perturbed regulatory networks were then compared with the corresponding experimental data.

**Table 2.** Data used for model calibration, corresponding simulation results, and parameters applied to replicate the experimental conditions.

| Experimental Data and Observations | Model Simulations | Adjusted Parameters |
| --- | --- | --- |
| Flow cytometry shows more E2-exposed cells in S phase than controls (Fig. 1D in Obiorah et al.<sup>31</sup>) | When E2 is ON, the death score drops from 0.018 to 0.002 but rises to 0.99 once the apoptosis pathway activates. | E2 = 1 |
| Extrinsic apoptotic genes (FAS, TNFα) are expressed 12 hours after intrinsic genes (BIM) (Figs. 4A-B in Lui et al.<sup>32</sup>). | With E2 ON, the extrinsic pathway (CASP8) is activated about 12.43 hours ($\pm 4.75$ hrs) after the intrinsic pathway (BIM). | |
| E2 induces the PERK cascade, and BAX is activated at 96 hours (Figs. 1B & 5A in Sengupta et al.<sup>21</sup>) | Also, E2 triggers the PERK/p-eIF2α/CHOP cascade, inducing intrinsic apoptosis (BIM) around 110.89 hours ($\pm 7.54$ hrs) | |
| Salubrinal induces intrinsic apoptosis, indicated by increased phosphorylated BAX (Fig. 5B in Sengupta et al.<sup>21</sup>). | Salubrinal triggers intrinsic apoptosis by enhancing E2-induced stress (death score = 0.99 when PP1 is OFF). | $PP1 = 0$ |
| Annexin V levels are lower in SRC siRNA samples than in controls (Fig. 2C in Obiorah et al.<sup>31</sup>) | E2-induced intrinsic apoptosis is blocked (BIM & BAX = 0) when SRC is suppressed due to inhibition of p-eIF2α. | $SRC = 0$<br>$E2 = 1$ |
| Everolimus increases the proportion of cells with autophagosomes in MCF7:5C compared to control (Fig. 8B in Lewis et al.<sup>20</sup>) | ATG14- BECN1, an autophagosome marker, is activated after mTORC1 inhibition, with a death score of 0.11. | $mTORC1 = 0$ |
| Everolimus-induced autophagosomes are reduced by Chloroquine (Fig. 8C in Lewis et al.<sup>20</sup>). | The combination of Chloroquine and an mTORC1 inhibitor shows a synergistic effect, with a Highest Single Agent (HSA)* score of 0.38 | $mTORC1 = 0$<br>$BECN1 = 0$ |
| mRNA levels of the extrinsic apoptotic gene such as TNFα increase after treatment with both E2 and the JNK inhibitor SP600125 (Fig. 3B in Fan et al.<sup>23</sup>). | The extrinsic pathway (TNFα-FADD-CASP8) is activated by the combination of SP600125 (a JNK inhibitor) and E2. | $E2 = 1$<br>$MAPK8 = 0$ |
| E2-induced extrinsic apoptosis (TNFα cascade) is blocked by JSH-23 (TNFα=0). TNFα mRNA levels are reduced by the combination of E2 and JSH-23 (Fig. 1H in Fan et al.<sup>22</sup>). | E2-induced extrinsic apoptosis (TNFα cascade) is blocked by JSH-23, reducing TNFα to zero. | $E2 = 1$<br>$NF\kappa B = 0$ |

\* HSA (Highest Single Agent) model evaluates drug combinations by comparing the observed combination effect, $E_{AB}$ , to the stronger of the two single-agent effects, defined as $\max(E_A, E_B)$ . A combination is considered synergistic if $E_{AB} > \max(E_A, E_B)$ . If $E_{AB} < \max(E_A, E_B)$ , the interaction is classified as antagonistic. If $E_{AB} = \max(E_A, E_B)$ , the combination is considered non-interactive (or equivalent to the best single agent effect), often described as additive under the HSA framework.

**Table 3.** Initial states of model variables, as inferred from experimental data obtained using the MCF7:5C cell line.

| Boolean variable | Initial state | Description | Reference |
| --- | --- | --- | --- |
| *E2* | 0 | MCF7:5C cells survive independently of estrogen (E2) | Fig. 3 in Escher et al.<sup>33</sup> |
| *ESR1* | 1 | ESR1 is highly expressed in MCF7:5C compared with MCF7 and T47D | Fig. 1A in Soung et al.<sup>34</sup> |
| *PERK* | 0 | PERK, eIF2α, ATF4, and CHOP are slightly phosphorylated in unperturbed MCF7:5C | Fig. 1B in Sengupta et al.<sup>21</sup> |
| *eIF2α* | 0 |  |  |
| *ATF4* | 0 |  |  |
| *CHOP* | 0 |  |  |
| *PP1* | 1 | PP1 remains active in both unperturbed and perturbed MCF7:5C | Fig. 2B in Sengupta et al.<sup>21</sup> |
| *AKT* | 1 | Phosphorylated AKT remains active in both unperturbed and perturbed MCF7:5C | Fig. 1A in Obiorah et al.<sup>31</sup> |
| *NFκB* | 0 | MCF7:5C cells show reduced NFκB expression relative to MCF7 and MCF7:2A | Fig. 1A in Fan et al.<sup>22</sup> |
| *STAT3* | 0 | E2 increases STAT3 activity at 72 hours compared to the vehicle in the western blot experiment | Fig. 6A in Fan et al.<sup>22</sup> |
| *CEBPB* | 0 | Baseline CEBPB expression in control is lower than in MCF7:5C cells following E2 treatment | Fig. 3A in Fan et al.<sup>22</sup> |
| *ERK* | 1 | Assuming ERK activation is driven by SRC presence in MCF7:5C | Fig. 1A in Obiorah et al.<sup>31</sup> |
| *MYC* | 1 | Assuming MYC activation is due to CCND1 presence | Fig. 2C in Lewis et al.<sup>20</sup> |
| *CCND1* | 1 | Phosphorylated CCND1 levels are high at 72 hours in unperturbed cells, comparable to those in Everolimus-treated cells | Fig. 2C in Lewis et al.<sup>20</sup> |
| *PI3K* | 1 | Assuming PI3K is active initially, given that AKT is active | Fig. 1A in Obiorah et al.<sup>31</sup> |
| *mTORC1* | 1 | mTOR is highly phosphorylated at the basal level in MCF7:5C | Fig. 4 in Lewis et al.<sup>20</sup> |
| *ATG14* | 0 | Assuming autophagy is inactive due to the low number of phagosomes observed in MCF7:5C | Fig. 8A in Lewis et al.<sup>20</sup> |
| *BECN1* | 0 |  |  |
| *SRC* | 1 | Baseline SRC phosphorylation in control is higher than in MCF7:5C cells after E2 treatment | Fig. 1A in Obiorah et al.<sup>31</sup> |
| *MAPK8* | 1 | Phosphorylated JNK levels are elevated in MCF-7:5C cells compared to MCF7 and MCF7:2A | Fig. 1A in Fan et al. <sup>23</sup> |
| *TNFα* | 0 | TNFα expression is lower in control than after E2 treatment | Fig. 4A in Fan et al. <sup>22</sup> |
| *TNFR1* | 0 | Assuming that TNFR1 is inactive due to the low levels of TNFα | Fig. 4A in Fan et al. <sup>22</sup> |
| *FADD* | 0 | Assuming that FADD is inactive due to the low levels of downstream proteins, specifically TNFα and TNFR1 | Fig. 4A in Fan et al. <sup>22</sup> |
| *CASP8* | 0 | The cleaved form of caspase-8 is not detected in MCF7:5C cells | Fig. 3B in Lewis et al. <sup>26</sup> |
| *BCL2* | 1 | BCL2 is highly phosphorylated in MCF7:5C | Fig. 5A in Sengupta et al. <sup>21</sup> |
| *BIM* | 0 | In unperturbed MCF7:5C cells, the phosphorylation levels of BIM and BAX are lower compared to those observed following E2 perturbation. | Fig. 5A in Sengupta et al. <sup>21</sup> |
| *BAX* | 0 |  | Fig. 5A in Sengupta et al. <sup>21</sup> |
| *CYCS* | 0 | Assuming that CYCS, CASP9, and CASP7 are inactive in MCF7:5C cells due to the low levels of cleaved CASP8, BIM, and BAX. Since CASP3 is truncated in MCF7:5C, with only detectable mRNA <sup>35</sup>, CASP7 is used as a substitute apoptosis executioner and as a proxy for CASP3/7 activity | Fig. 3B in Lewis et al. <sup>26</sup> |
| *CASP9* | 0 |  | Fig. 3B in Lewis et al. <sup>26</sup> |
| *CASP7* | 0 |  | Fig. 3B in Lewis et al. <sup>26</sup> |

### Setting initial states

We collected data from the literature to define the initial states of proteins in MCF7:5C cells. Table 3 summarizes the initial states of the Boolean variables for the corresponding proteins, the rules used to determine these values, and the literature sources from which the data for the initial states were derived. The following assumptions were applied to determine the initial activity state (Boolean value) of each protein: (i) the initial protein activity is set to 0 if the differential expression of the corresponding gene in MCF7:5C is lower than in other cancer cell lines, and set to 1 otherwise, (ii) the initial protein activity is set to 0 if the protein activity in control (untreated) MCF7:5C cells is lower than in drug-perturbed MCF7:5C cells; otherwise, it is set to 1, and (iii) the initial protein activity is set to 0 if either upstream regulators or downstream targets of the protein are known to have low activity; otherwise, it is set to 1. The death score was calculated using the initial protein states listed in Table 3 and Equation (9). The initial global loading state was derived from the methodology described in the Supplementary Text (S1 Text).

### Perturbation analysis

We applied the calibrated model to simulate single- and double-node perturbations within the cell death regulatory network to identify those that inhibit cell death in E2-treated MCF7:5C cells. These perturbations were simulated by modifying the parameters *ω*_*i*_ and *ω*_*ij*_ such that the variables describing protein activity are set to either 1 or 0 throughout the simulation, representing gain or loss of corresponding protein function. For example, a gain-of-function mutation in the AKT protein is modeled by setting all *ω*_AKT,j_ to 1 and *ω*_AKT_ to 10, while a loss-of-function mutation is modeled by setting *ω*_AKT,j_ to 0 and *ω*_AKT_ to −10 throughout the simulation.

To determine when the perturbed network reaches a steady state, we evaluated network state *S* using Shannon entropy, calculated over the time interval from 0 to *t*, with a maximum of 1000 iterations. The entropy is defined as:

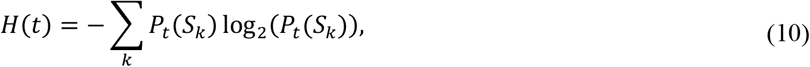

where 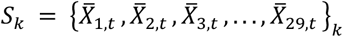 represents the *k*-th unique set of averaged protein activity values 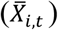 observed over the time interval from 0 to *t*, averaged across 30 simulation repeats. The probability *P*_*t*_(*S*_*k*_) was computed as the ratio of the number of times state *S*_*k*_ occurs to the total number of states observed within the time interval from 0 to *t*. As the system approaches a defined steady state, entropy gradually decreases and tends toward zero. However, since it is impractical to wait for entropy to reach exactly zero, we instead calculated the slope of entropy over time (within the 0 to 1000 time-step window). The time point, *T*_*ss*_, corresponding to the steepest negative slope (i.e., the fastest rate of entropy decreases) was identified as the point at which the system reaches a steady state.

Using this approach, we independently determined the steady state point, *T*_*ss*_, for different simulation scenarios. First, for MCF7:5C cells subjected to single- and double-node perturbations, *T*_*ss*_ was used as the point to introduce E2 treatment. This allowed us to identify perturbations that prevent E2-induced apoptosis, indicating potential resistant cell strains. Resistance was defined based on the inactivity of CASP7 at steady state (i.e., *CASP7* = 0 at *T*_*ss*_ in perturbed cells with E2 treatment), serving as a marker of failed apoptosis. Next, we applied a second round of single-node perturbations by activating or inactivating one more node, to identify additional targets capable of reinducing apoptosis in these resistant strains. Resistant strains in which CASP7 became active at steady state (*CASP7* = 1 at *T*_*ss*_ following the second-round perturbation) were classified as re-sensitized to apoptosis.

## Results

### The Boolean model accurately captures E2-induced apoptosis in MCF7:5C

The effect of E2 treatment on key cell death regulators and execution pathways in MCF7:5C cells was simulated using 30 asynchronous update repeats, with E2 activation (*E2* = 1) occurring at 50 hours (indicated by the red dashed line in Fig 2A). Before E2 treatment, MCF7:5C cells exhibit E2-independent survival, characterized by a low death score (green dashed line in Fig 2A). Upon E2 exposure, the death score initially decreases due to increased protein synthesis (*GL*, blue dashed line in Fig 2A). However, prolonged exposure to E2 leads to excessive protein synthesis, which activates the PERK pathway (brown dashed line in Fig 2A). This activation increases the level of phosphorylated eIF2α, ultimately resulting in a significant reduction in global protein synthesis. The behavior of these components is regulated by a negative feedback loop: GL activates PERK, which phosphorylates eIF2α that subsequently inhibits further protein synthesis (GL → PERK → p-eIF2α ⊣ GL; see Fig 1). In addition, phosphorylated EIF2α can activate the ATF4 and CHOP proteins, which in turn initiate the intrinsic apoptosis pathway via BAX activation (blue line in Fig 2B). Extrinsic apoptosis is also induced through the PERK → STAT3 → NFκB → TNFα signaling cascade following activation of the intrinsic pathway (red line in Fig 2B) ^22^. Beyond capturing the general characteristics of the E2 response, our model is specifically tuned to reproduce the experimental observations summarized in Table 2.

**Fig 2.**
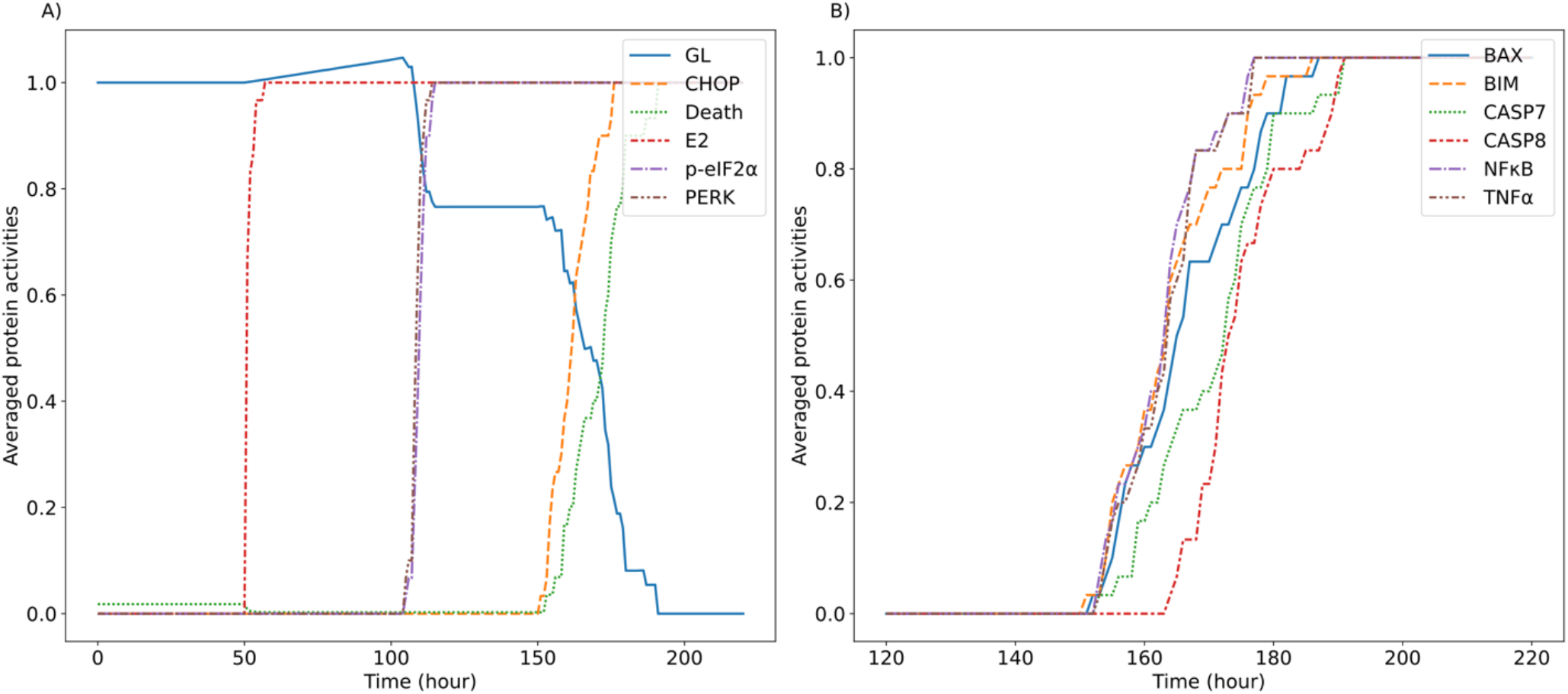
Dynamic behavior of key model components in MCF7:5C cells over 200 hours following E2 treatment at 50 hours. Average protein activities from 30 simulations involved in (A) PERK pathway components. (B) Intrinsic and extrinsic apoptosis pathways.

### Blocking PERK signaling via pathway component deletion mitigates E2 treatment effects in MCF7:5C cells

Perturbation analysis was performed to identify alterations that could block E2-induced apoptosis in MCF7:5C cells. To simulate the effect of single-gene mutations, each node in the signaling network shown in Fig 1 was individually forced to be constitutively activated (node variable = 1) or inactivated (node variable = 0). Fig 3 shows the resulting steady-state protein activity levels from these perturbations. To determine whether the cell survives a given perturbation, we recorded the states of CCND1, mTORC1, BAX, CASP7, BECLN1, and CASP8, as these proteins serve as key indicators contributing to the calculated death score (see, Equation 9).

**Fig 3.**
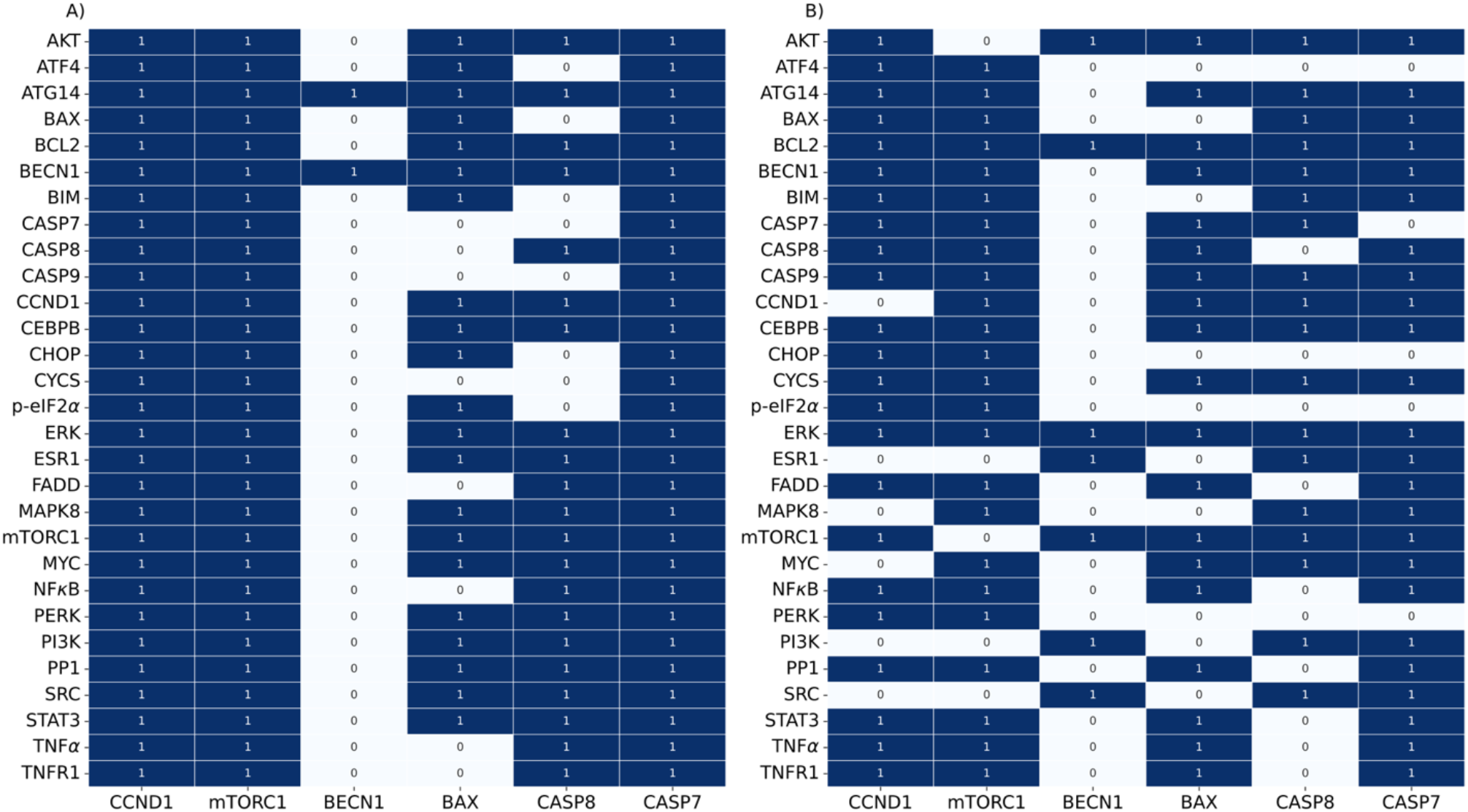
Steady-state protein activities are shown with readout proteins on the x-axis and perturbed proteins on the y-axis. Panel (A) represents perturbations where nodes were set to one (activated), while panel (B) shows perturbations where nodes were set to zero (inactivated). Values of 1 and 0 correspond to the average protein activity levels from 30 simulation repeats.

The PERK pathway mediates E2-induced apoptosis through both extrinsic ^22^ and intrinsic mechanisms ^21^ in MCF7:5C cells. Our results (Fig 3B) predict that deletions in critical PERK pathway components (PERK, p-eIF2α, ATF4, and CHOP) enable MCF7:5C cells to evade E2-induced cytotoxicity by simultaneously suppressing both apoptotic pathways. This survival advantage is further supported by the sustained activation of pro-survival signaling components, including mTORC1 and CCND1. Additionally, the results (Fig 3B) show that direct inhibition of the effector caspase CASP7 contributes to the loss of apoptotic function in MCF7:5C cells, a mechanism also reported in other cancer types ^34^. This finding explains why the model classifies CASP7-deficient cells as resistant to E2 therapy. Overall, the following features were identified as resistant to E2 treatment: *PERK*^−^, p-*eIF2α*^−^, *ATF4*^−^, *CHOP*^−^, and *CASP7*^−^, where the superscript minus sign denotes deficiency or deletion of the corresponding gene or protein.

For all perturbations that block E2-induced apoptosis in MCF7:5C cells, we investigated additional single-node interventions that enable these cells to overcome resistance to E2 treatment. We found that inhibition of Protein Phosphatase 1 (PP1) in *PERK*^−^ cells restores activation of the intrinsic apoptosis pathway, see Table 4. This occurs because reduced PP1 activity limits dephosphorylation of eIF2α, thereby activating the p-eIF2α → ATF4 → CHOP → BIM signaling cascade ^21^, whereas activation of PERK in this cell can activate both intrinsic and extrinsic apoptosis pathway ^31^. Furthermore, the intrinsic apoptosis was induced in *PERK*^−^, *p*−*eIF*2α^−^, *CHOP*^−^, and *ATF*4^−^ cells by the direct activation of gene downstream of the deletion. For example, direct activation of p-eIF2α/ATF4/CHOP pathway promoted intrinsic apoptosis in *PERK*^−^ and *p*−*eIF*2α^−^ cells, while *CHOP*^−^, and *ATF*4^−^ cells are limited to the activation of CHOP, BAX, and BIM.

**Table 4.** Single-node interventions that restore E2-induced apoptosis in resistant MCF7:5C cells.

| E2 resistant strain | Perturbation type | List of perturbed nodes that induce apoptosis in resistant strains |  |  |
| --- | --- | --- | --- | --- |
|  |  | Intrinsic Apoptosis | Extrinsic Apoptosis | Other |
| *PERK*<sup>-</sup> | Protein inhibition | PP1 | SRC, PI3K, ESR1, MAPK8 | - |
| *p - eIF2α*<sup>-</sup> |  | - | SRC, PI3K, ESR1, MAPK8, CEBPB | - |
| *ATF4*<sup>-</sup> |  | - | SRC, PI3K, ESR1, MAPK8, CEBPB | - |
| *CHOP*<sup>-</sup> |  | - | SRC, PI3K, ESR1, MAPK8, CEBPB | - |
| *PERK*<sup>-</sup> | Protein activation | ATF4, BAX, BIM, CHOP, p-eIF2α | FADD, TNFα, TNFR1, NFκB, CASP8 | CYCS*, CASP7*, CASP9*, PERK** |
| *p - eIF2α*<sup>-</sup> | | CHOP, BAX, BIM, p-eIF2α, ATF4 | FADD, TNFα, TNFR1, NFκB, CASP8 | CASP9*, CASP7*, CYCS* |
| *ATF4*<sup>-</sup> | | CHOP, BAX, BIM, ATF4 | FADD, TNFα, TNFR1, NFκB, CASP8 | CASP9*, CASP7*, CYCS* |
| *CHOP*<sup>-</sup> | | CHOP, BAX, BIM | FADD, TNFα, TNFR1, NFκB, CASP8 | CASP9*, CASP7*, CYCS* |

\*Directly activating downstream of apoptosis pathway.
\*\*Activating both extrinsic and intrinsic apoptosis

Extrinsic apoptosis was induced by activation of key components of the TNF signaling pathway, including FADD, TNFα, TNFR1, CASP8, and NFκB or by inhibition of SRC, PI3K, ESR1, CEBPB, MAPK8. This is summarized as diagram, which illustrate how the extrinsic apoptosis through TNFα → TNFR1 → FADD → CASP8 axis occurred in the mutant strain in Table 4 (red box in Fig.4A), either by relief of negative regulation of downstream proteins of TNFα axis (blue box in Fig.4A) or through activation of signaling components downstream of TNFα (yellow box in Fig.4A). To illustrate this mechanism dynamically, Fig 4B shows an example of the time courses of key protein activities during extrinsic apoptosis by reliving the negative regulation of TNFα axis by MAPK8 inhibition in the presence of E2 in the *PERK*^−^ strain., which subsequently mediate the TNFα → TNFR1 → FADD → CASP8 axis.

**Fig 4.**
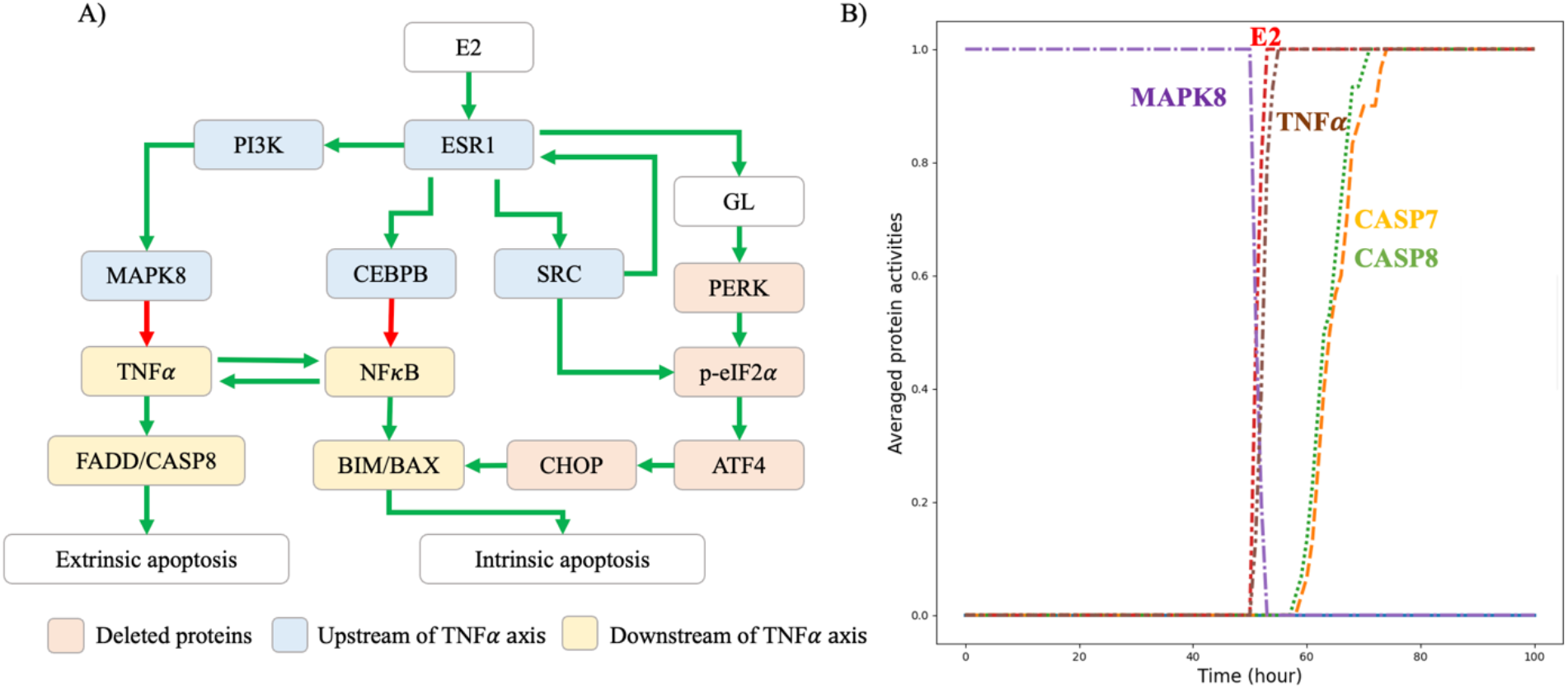
(A) Signal rerouting achieved by inhibition of target proteins (purple rectangles) redirects E2-induced signaling away from pathways blocked by gene deletions (red rectangles). (B) Time courses of key protein activities, averaged over 30 simulation repeats, during induction of extrinsic apoptosis by MAPK8 inhibition (MAPK8 = 0) in the presence of estradiol (E2 = 1) in the *PERK*^−^ strain.

Overall, perturbation analysis revealed four distinct routes leading to MCF7:5C cell death: (i) activation of the intrinsic apoptotic pathway via PERK axis, (ii) activation of the extrinsic apoptotic pathway via either inhibiting upstream or activating downstream of TNFα genes, (iii) concurrent activation of both pathways via upregulating PERK protein, and (iv) direct activation of apoptotic effector proteins (CYCS, CASP9, and CASP7).

### Predicting double-node perturbations blocking E2-induced apoptosis

To explore more complex E2-resistant strains of MCF7:5C cells, we simulated possible double-node perturbations in the apoptotic regulatory network shown in Fig 1. For each pair of nodes X and Y, four perturbation combinations were simulated: *X*^+^*Y*^+^, *X*^+^*Y*^−^, *X*^−^*Y*^+^, and *X*^−^*Y*^−^, where the superscript minus sign indicates node inactivation and the superscript plus sign indicates node activation.

To focus on genes that frequently acquire mutations in cancer cells, we analyzed mutation data from the DepMap database and quantified the frequency of gene-damaging mutations across cancer cell lines for 29 genes involved in the apoptotic regulatory network shown in Fig 1. The 15 genes with the highest mutation frequencies were then selected for systematic evaluation of all possible double-gene perturbations (Fig 5A). In each group (i.e., *X*^+^*Y*^+^, *X*^+^*Y*^−^, *X*^−^*Y*^+^, and *X*^−^*Y*^−^), there are fifteen selected genes, resulting in 105 pairwise combinations per group, calculated using the binomial coefficient 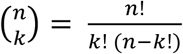 with n=15 and k = 2, to simulate double-mutant strains under E2 treatment. Overall, double-node perturbation analysis identified 58 E2-resistant MCF7:5C strains, with PERK among the most frequently perturbed genes in these strains (Fig 5B). This finding is consistent with previous reports identifying PERK as a key regulator of E2-induced signaling that activates both intrinsic and extrinsic apoptotic pathways in MCF7:5C cells ^22^.

**Fig 5.**
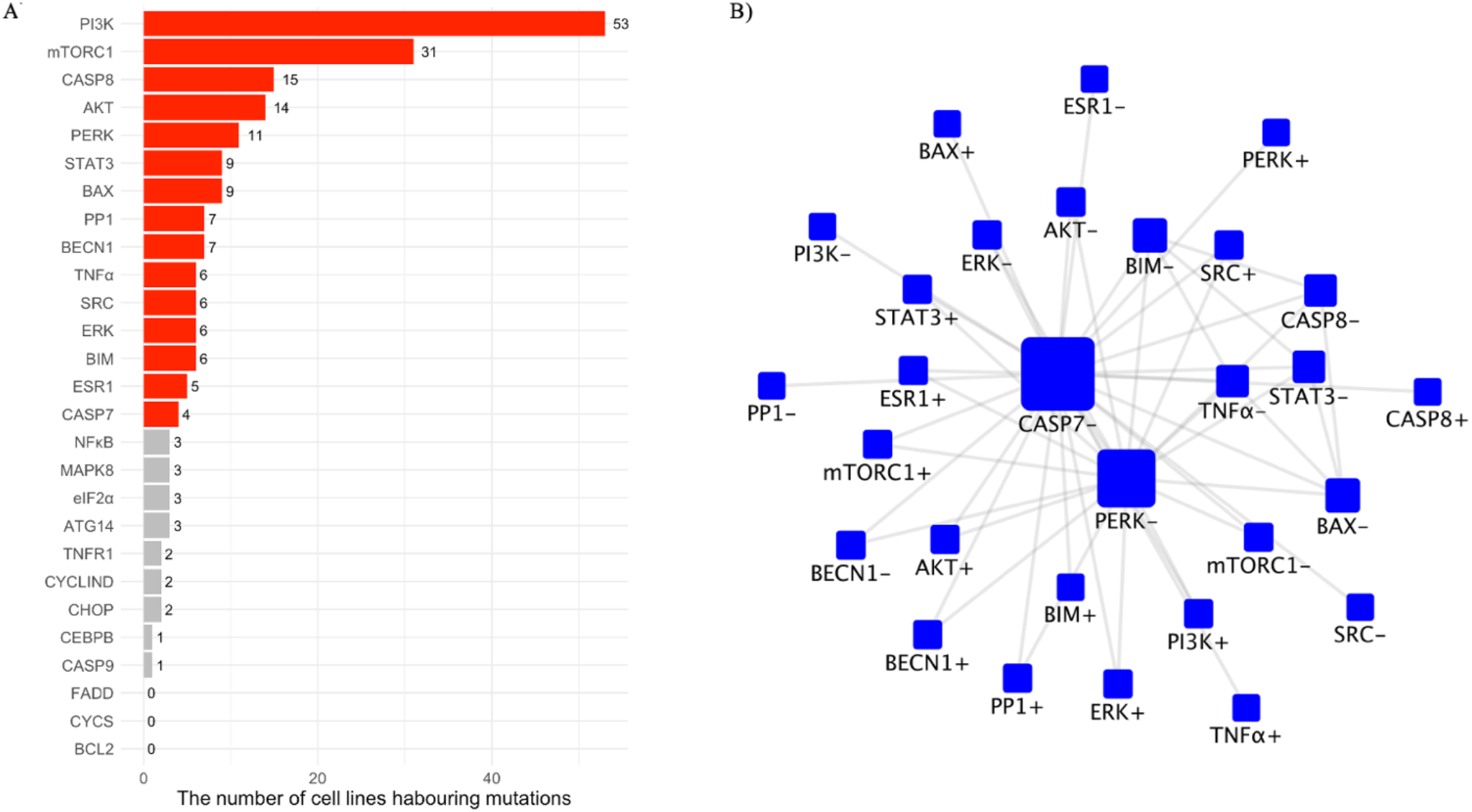
(A) Frequencies of gene-damaging mutations across 1,788 human cancer cell lines from 29 lineages based on analysis of DepMap Public 24Q2. Red bars denote the 15 genes with the highest mutation frequencies among genes in the apoptotic regulatory network (red color). (B) Network summarizing the involvement of perturbed nodes in 58 strains exhibiting E2 resistance. Node size reflects node degree, indicating how frequently each gene appears in double-node perturbations associated with E2 resistance.

Among the 58 E2-resistant strains, 28 involved CASP7 inactivation. In these cases, apoptosis can be restored only via direct reactivation of CASP7. In the remaining 30 strains, additional single-node perturbations were systematically evaluated to identify conditions that restore E2-induced apoptosis. Fig 6 summarizes model predictions for 506 possible single-node interventions that restore E2-induced cell death across 30 E2-resistant strains with double-acquired mutations. The interventions are grouped into four distinct modes: (i) direct activation of CASP7 (orange tiles), (ii) activation of the extrinsic apoptotic pathway (red tiles), (iii) activation of the intrinsic apoptotic pathway (blue tiles), and (iv) simultaneous activation of both intrinsic and extrinsic apoptotic pathways (grey tiles).

**Fig 6.**
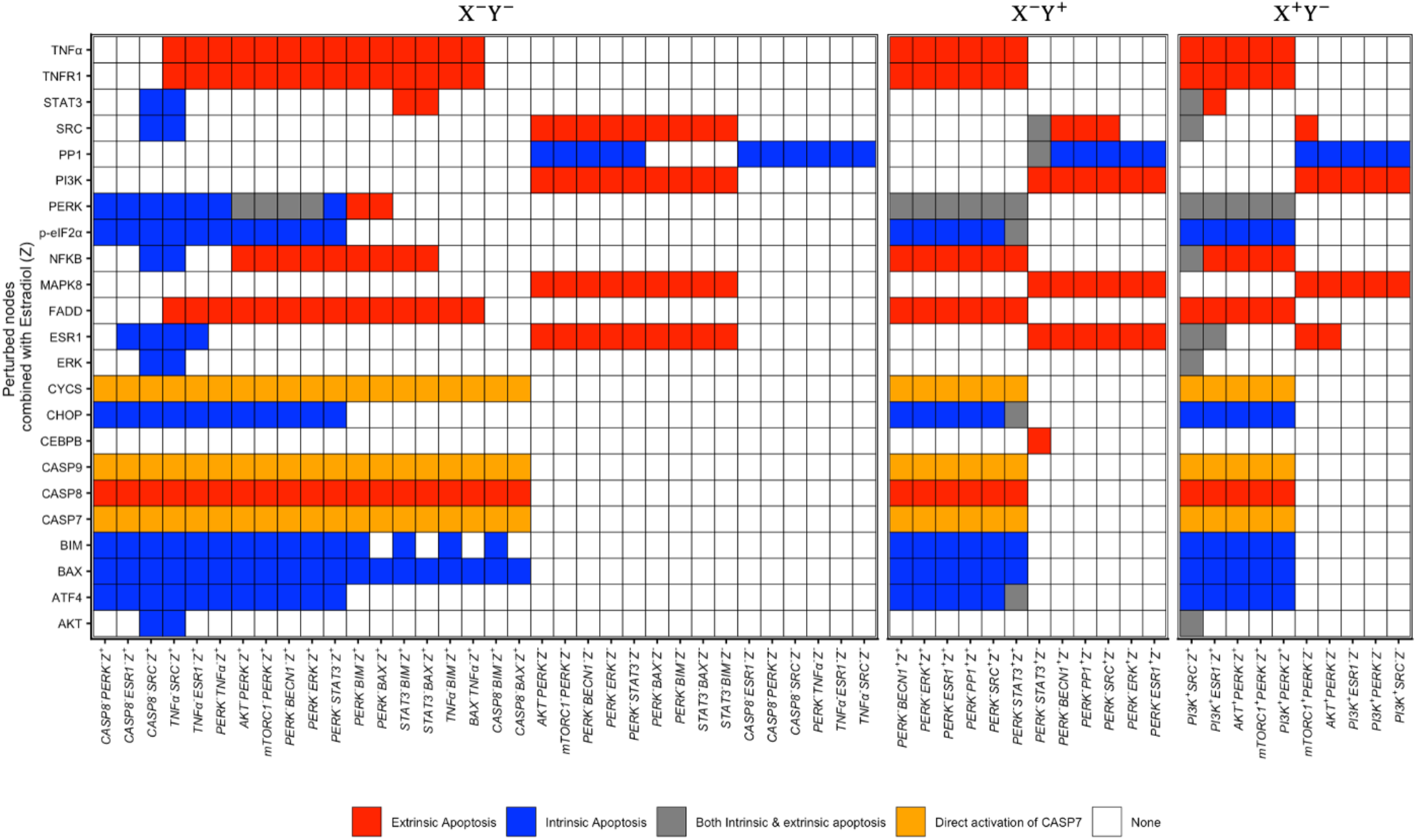
Model-predicted single-node perturbations capable of restoring E2-induced cell death in 24 E2-resistant strains. Each column represents a resistant strain combined with a perturbation of gene *Z* (y-axis). Superscript minus (^−^) and plus (^+^) indicate node inactivation and activation, respectively. E2-resistant strains are grouped as follows: Group 1 (*X*^−^*Y*^−^, left panel), in which both genes are inactive; Group 2 (*X*^−^*Y*^+^, middle panel), in which the first gene is inactive and the second is active; and Group 3 (*X*^+^*Y*^−^, right panel), in which the first gene is active and the second is inactive. Grid tiles are colored according to the observed mode of cell death, as indicated in the figure legend.

In the first mode, direct activation of apoptotic effector proteins, including CASP7, CASP9, and CYCS, restores apoptosis in all resistant strains from groups 1–3 (orange tiles in Fig 6), independent of whether intrinsic or extrinsic pathways are engaged under E2 treatment, consistent with the role of CASP7 as an executioner caspase ^34^.

Extrinsic apoptosis (the second death mode) was reactivated in most E2-resistant strains across Groups 1–3 (red tiles in Fig 6) upon activation of components of the NFκB → TNFα → TNFR1 → FADD → CASP8 pathway. In contrast, strains harboring loss-of-function mutations in CASP8, (e.g., *CASP*8^−^ *PERK*^−^, *CASP*8^−^ *BIM*^−^ and *CASP*8^−^ *BAX*^−^) failed to restore extrinsic apoptosis unless CASP8 was directly reactivated. Inactivation of PI3K, MAPK8, ESR1, or SRC restored E2-induced extrinsic apoptosis by relieving the negative regulation of TNFα and NFκB in the double-mutant strains lacking PERK. The model further predicted that downregulation of CEBPB (*CEBPB* = 0) rescues E2-induced extrinsic apoptosis in the *PERK*^−^ *STAT*3^+^ strain, whereas upregulation of STAT3 (*STAT3* = 1) restores E2-induced extrinsic apoptosis in *STAT*3^−^ *BAX*^−^ and *STAT*3^−^ *BIM*^−^ strains through activation of the NFκB → TNFα → TNFR1 signaling axis.

Intrinsic apoptosis (the third death mode) was reinduced in most E2-resistant strains across Groups 1–3 (blue tiles in Fig 6) following direct upregulation of the death executors BAX and BIM, or activation of the PERK pathway through upregulation of p-eIF2α, ATF4, or CHOP, or through downregulation of PP1. E2-resistant strains lacking functional death executors (e.g., *STAT*3^−^ *BAX*^−^ and *CASP*8^−^ *BIM*^−^ in Group 1) underwent E2-induced apoptosis only when BAX and BIM were directly reactivated. In addition to direct activation, our model predicts that STAT3 activation can induce intrinsic apoptosis through the NFKB → BIM → BAX pathway in strains lacking extrinsic apoptosis executors (e.g., *TNFα*^−^ *SRC*^−^ and *CASP8*^−^ *SRC*^−^). Furthermore, upregulation of AKT in these strains was predicted to promote intrinsic apoptosis by activating ESR1-mediated signaling and subsequently increasing global protein synthesis. This increase in protein synthesis is expected to trigger intrinsic apoptosis through activation of the PERK axis.

Both E2-induced extrinsic and intrinsic apoptosis signaling branches (the fourth death mode) were reactivated in E2-resistant strains when PERK was constitutively active (*PERK* = 1), as indicated by the grey tiles. Inactivation of SRC (*SRC* = 0) promoted E2-induced intrinsic apoptosis in the *PERK*^−^ *STAT*3^+^ strain by activating BIM and BAX through downregulation of ERK and AKT, and also induced extrinsic apoptosis by relieving negative regulation of NFκB via the SRC → ESR1 → CEBPB ⊣ NFκB pathway. Interestingly, in the *PI*3*K*^+^ *SRC*^−^ strain, AKT upregulation could trigger both intrinsic and extrinsic apoptosis through the PERK pathway by increasing global protein synthesis. Inactivation of PP1 (*PP1* = 0) similarly promoted both E2-induced intrinsic and extrinsic apoptosis in the *PERK*^−^ *STAT*3^+^ strain by decreasing dephosphorylation of p-eIF2α, thereby enhancing CHOP activity and activating both the intrinsic CHOP → BIM → BAX axis and the extrinsic CHOP → NFκB → TNFα → TNFR1 → FADD → CASP8 axis. Comparable effects were observed upon direct activation of PERK, p-eIF2α, ATF4, or CHOP.

## Discussion

While E2-based treatment has demonstrated clinical benefit in some postmenopausal women with aromatase inhibitor–resistant breast cancer ^18^, its therapeutic efficacy can be compromised by acquired mutations that emerge under treatment pressure. In this study, we developed a stochastic continuous-time Boolean model of the death-signaling network in the MCF7:5C breast cancer cell line to simulate single- and double-gene acquired mutations and to identify complementary interventions capable of restoring E2-induced apoptosis. The model accurately recapitulated quantitative time-course data for E2–induced responses of key cell death regulators, including intrinsic and extrinsic apoptotic effectors and upstream signaling components (Fig 1 and Table 2). This analysis helps to explain how cancer cells can bypass action of the treatment and how targeted interventions can restore cell sensitivity to therapy.

The model was used to explore mechanisms underlying E2 resistance in MCF7:5C cells and to identify potential strategies for restoring E2-induced cell death. The simulations underscore the central role of the PERK pathway in mediating E2-induced apoptosis, as loss-of-function alterations in PERK pathway components conferred resistance to E2 treatment. These findings highlight the PERK pathway as a key determinant of E2 responsiveness and a potential therapeutic target for overcoming E2 resistance. Notably, several model predictions align with prior experimental observations, as inhibition of SRC ^30^, MAPK8 ^23^, and NFκB ^22^ was predicted to restore E2-induced apoptosis in MCF7:5C cells with impaired PERK signaling. Beyond these validated targets, the model also identified multiple novel interventions. In particular, inhibition of PI3K, ESR1, or CEBPB was predicted to restore E2-induced extrinsic apoptosis through enhanced activation of downstream TNFα signaling, suggesting alternative therapeutic routes for overcoming E2 resistance when PERK signaling is compromised.

The analysis was further extended to capture more complex resistance phenotypes arising from combined gain- and loss-of-function alterations within the apoptotic regulatory network. Consistent with findings from single-gene perturbations, disruption of PERK signaling frequently emerged as a critical contributor to resistance in double-gene alteration scenarios, reinforcing its pivotal role in E2-induced cell death. Simulations revealed multiple targets capable of reactivating apoptotic signaling in MCF7:5C cells harboring dual alterations; however, therapeutic efficacy was highly context dependent. For instance, while PERK activation effectively induced both intrinsic and extrinsic apoptosis in certain double-mutant strains, it failed to restore cell death in strains with concurrent impairment of PERK and key apoptotic effectors such as BIM, BAX, or CASP8.

The observed heterogeneity in E2 responsiveness across distinct mutational backgrounds underscores the strain- and context-dependent nature of responses to specific therapies. These findings emphasize the importance of incorporating individualized genomic profiles when designing E2-based combination therapies for menopausal women, with the goal of maximizing therapeutic efficacy while minimizing resistance ^36^.

The present model successfully captures both wild-type and E2-resistant behaviors of the MCF7:5C cell line, but it is limited by its exclusive focus on a single mode of programmed cell death, namely apoptosis. Multiple additional forms of programmed cell death—including necroptosis, pyroptosis, ferroptosis, and immunogenic cell death—have been shown to play important roles in breast cancer cell fate determination and therapeutic response ^37–40^. Incorporating these alternative death pathways is essential for achieving a more comprehensive understanding of cell fate decision-making. Such extensions were not pursued in the current study due to the limited availability of experimental data characterizing drug perturbations and death pathway activation in MCF7:5C cells.

As more high-resolution experimental datasets become available, a key future direction will be to integrate crosstalk among multiple programmed cell death modalities into the modeling framework. Such an expanded model would enable systematic identification of novel regulators that govern cell death across distinct modes and facilitate the discovery of therapeutic targets capable of redirecting resistant cancer cells toward alternative, non-apoptotic death pathways in the presence of acquired mutations.

## Acknowledgments

K.T. acknowledges the Petchra Pra Jom Klao Ph.D. Research Scholarship (KMUTT – NSTDA) from King Mongkut’s University of Technology Thonburi (No: 103/2563).

## Data Availability

The model is available on https://github.com/Ktaoma/Stochastic-Boolean-Model-of-Death-Signaling-in-MCF7-5C-Predicts-Cell-Death-Inducers-and-Inhibitors

## Contributions

K.T., T.L., S.S., Y.W., R.C., and P.K. conceived the study. K.T., T.L., and P.K. developed the stochastic Boolean framework. K.T. developed the Python code, performed the simulations, and prepared the figures. K.T. and P.K. analyzed and interpreted the simulation results. All authors contributed to writing and revising the manuscript.

Disclaimer: Surojeet Sengupta is currently employed at Center for Scientific Review, National Institutes of Health. This article was prepared while Surojeet Sengupta was employed at The Hormel Institute. The opinions expressed in this article are the author’s own and do not reflect the view of the National Institutes of Health, the Department of Health and Human Services, or the United States government.

## Supplementary information

S1 Text: The derivation of global protein synthesis (λ) and reduction rate (β) parameters from Western blot data.

Given the slope equation of global protein synthesis, 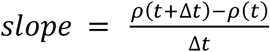, we can integrate the global loading as follows:

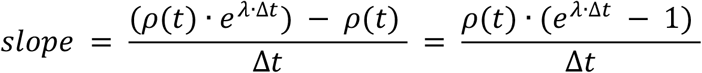

if *λ* · Δ*t* is small, *e*^*λ*·Δ*t*^ can be approximated as 1 + (*λ* · Δ*t*). Thus, the slope becomes:

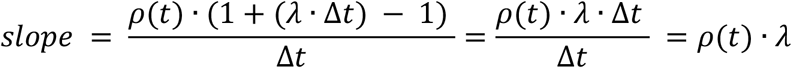

Therefore, *λ* can be rearranged as:

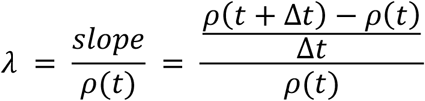

To calculate *λ*, we digitized the experimental data from Figure 1A of Surojeet et al. 2019 (PMID: 30655322) using ImageJ software (Version 1.54m) into the following table:

|  | Intensity |
| --- | --- |
| Vehicle | 107.653 |
| 4 hrs. | 115.026 |
| 24 hrs. | 124.841 |
| 48 hrs. | 119.328 |
| 72 hrs. | 86.887 |
| 96 hrs. | 85.932 |

Then, the global protein synthesis rate was calculated as:

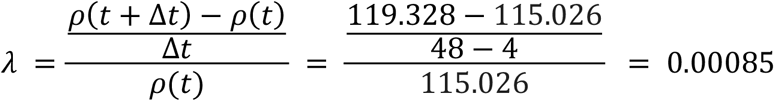

The reduction rate *β* was calculated as:

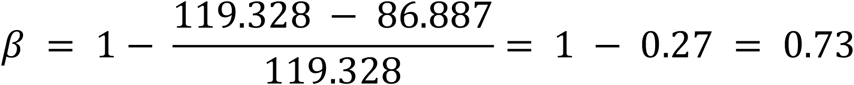

**S1 Table:** List of interactions in the signaling network associated with E2-induced apoptosis in MCF7-5C cells and the sources used to identify and curate these interactions.

| source | target | type | reference | Note |
| --- | --- | --- | --- | --- |
| PI3KCA | AKT | Activation | Signor Database | direct activation |
| SRC | AKT | Activation | Signor Database | direct activation |
| p-eIF2α | ATF4 | Activation | PMID: 30655322 | direct activation |
| mTORC1 | ATG14 | Inhibition | Signor Database | Indirect inhibiton through ULK1<br>(mTORC1 $\dashv$ ULK1 → ATG14) |
| BCL2 | BAX | Inhibition | Signor Database | direct activation |
| BIM | BAX | Activation | Signor Database | direct activation |
| CASP8 | BAX | Activation | Signor Database | indirect activation through BID<br>(CASP8 → BID → BAX) |
| AKT | BCL2 | Activation | Signor Database | indirect activation through GSK3B<br>(AKT $\dashv$ GSK3B $\dashv$ BCL2) |
| BIM | BCL2 | Inhibition | Signor Database | direct activation |
| ERK | BCL2 | Activation | Signor Database | direct activation |
| ATF4 | BECN1 | Activation | PMID: 34168552 | direct activation |
| ATG14 | BECN1 | Activation | Signor Database | indirect activation through PIK3C3<br>(ATG14 → PIK3C3 → BELIN1) |
| BCL2 | BECN1 | Inhibition | Signor Database | direct activation |
| AKT | BIM | Inhibition | Signor Database | direct activation |
| ERK | BIM | Inhibition | Signor Database | direct activation |
| NFκB | BIM | Activation | PMID: 29531812 | direct activation |
| CHOP | BIM | Activation | PMID: 30655322 | direct activation |
| CASP9 | CASP3 | Activation | Signor Database | direct activation |
| CASP7 | CASP7 | Activation | Signor Database | direct activation |
| CASP8 | CASP7 | Activation | Signor Database | direct activation |
| FADD | CASP8 | Activation | Signor Database | direct activation |
| AKT | CASP9 | Inhibition | Signor Database | direct activation |
| CASP8 | CASP9 | Activation | Signor Database | direct activation |
| CYCS | CASP9 | Activation | Signor Database | direct activation |
| AKT | CCND1 | Activation | Signor Database | indirect activation through GSK3B<br>(AKT → G2K3B → CCND1) |
| MYC | CCND1 | Activation | Signor Database | direct activation |
| ESR1 | CEBPB | Activation | PMID: 29531812 | direct activation |
| ATF4 | CHOP | Activation | PMID: 30655322 | direct activation |
| BAX | Death | Activation | Signor Database | direct activation |
| CASP7 | Death | Activation | Signor Database | direct activation |
| CASP8 | Death | Activation | Signor Database | direct activation |
| PERK | p-eIF2α | Activation | PMID: 30655322 | direct activation |
| PP1 | p-eIF2α | Inhibition | PMID: 30655322 | direct activation |
| SRC | p-eIF2α | Activation | PMID: 23704208 | direct activation |
| SRC | ERK | Activation | PMID: 23704208 | direct activation |
| AKT | ESR1 | Activation | Signor Database | direct activation |
| E2 | ESR1 | Activation | Signor Database | direct activation |
| ERK | ESR1 | Activation | Signor Database | direct activation |
| SRC | ESR1 | Activation | Signor Database | direct activation |
| TNFR1 | FADD | Activation | Signor Database | direct activation |
| p-eIF2α | GL | Inhibition | PMID: 30655322 | direct activation |
| ESR1 | GL | Activation | PMID: 30655322 | direct activation |
| E2 | GL | Activation | PMID: 30655322 | direct activation |
| PI3K | MAPK8 | Activation | Signor Database | indirect activation through RAC1<br>(PI3K → RAC1 → MAPK8) |
| AKT | mTORC1 | Activation | Signor Database | direct activation |
| ERK | MYC | Activation | Signor Database | direct activation |
| ESR1 | MYC | Activation | Signor Database | direct activation |
| MAPK8 | MYC | Activation | Signor Database | direct activation |
| mTORC1 | MYC | Activation | Signor Database | direct activation |
| CEBPB | NFκB | Inhibition | PMID: 29531812 | direct activation |
| CHOP | NFκB | Activation | PMID: 29531812 | direct activation |
| STAT3 | NFκB | Activation | PMID: 29531812 | direct activation |
| GL | NFκB | Activation | PMID: 30655322 | direct activation |
| PERK | NFκB | Activation | PMID: 29386355 | direct activation |
| ESR1 | PI3K | Activation | Signor Database | direct activation |
| ATF4 | PP1 | Activation | PMID: 30655322 | direct activation |
| CCND1 | SRC | Activation | Signor Database | indirect activation through NOTCH1<br>(CCND1 → NOTCH1/ERBB2/PTK2 → SRC) |
| ESR1 | SRC | Activation | PMID: 39413983 | direct activation |
| MAPK8 | STAT3 | Activation | Signor Database | direct activation |
| PERK | STAT3 | Activation | PMID: 29531812 | direct activation |
| BAX | Survival | Activation | Signor Database | direct activation |
| BECN1 | Death | Inhibition | Signor Database | direct activation |
| CCND1 | Death | Inhibition | Signor Database | direct activation |
| mTORC1 | Death | Inhibition | Signor Database | direct activation |
| MAPK8 | TNFα | Inhibition | PMID: 26116171 | direct activation |
| NFκB | TNFα | Activation | PMID: 29531812 | direct activation |
| TNFα | TNFR1 | Activation | Signor Database | direct activation |

**S2 Table:**
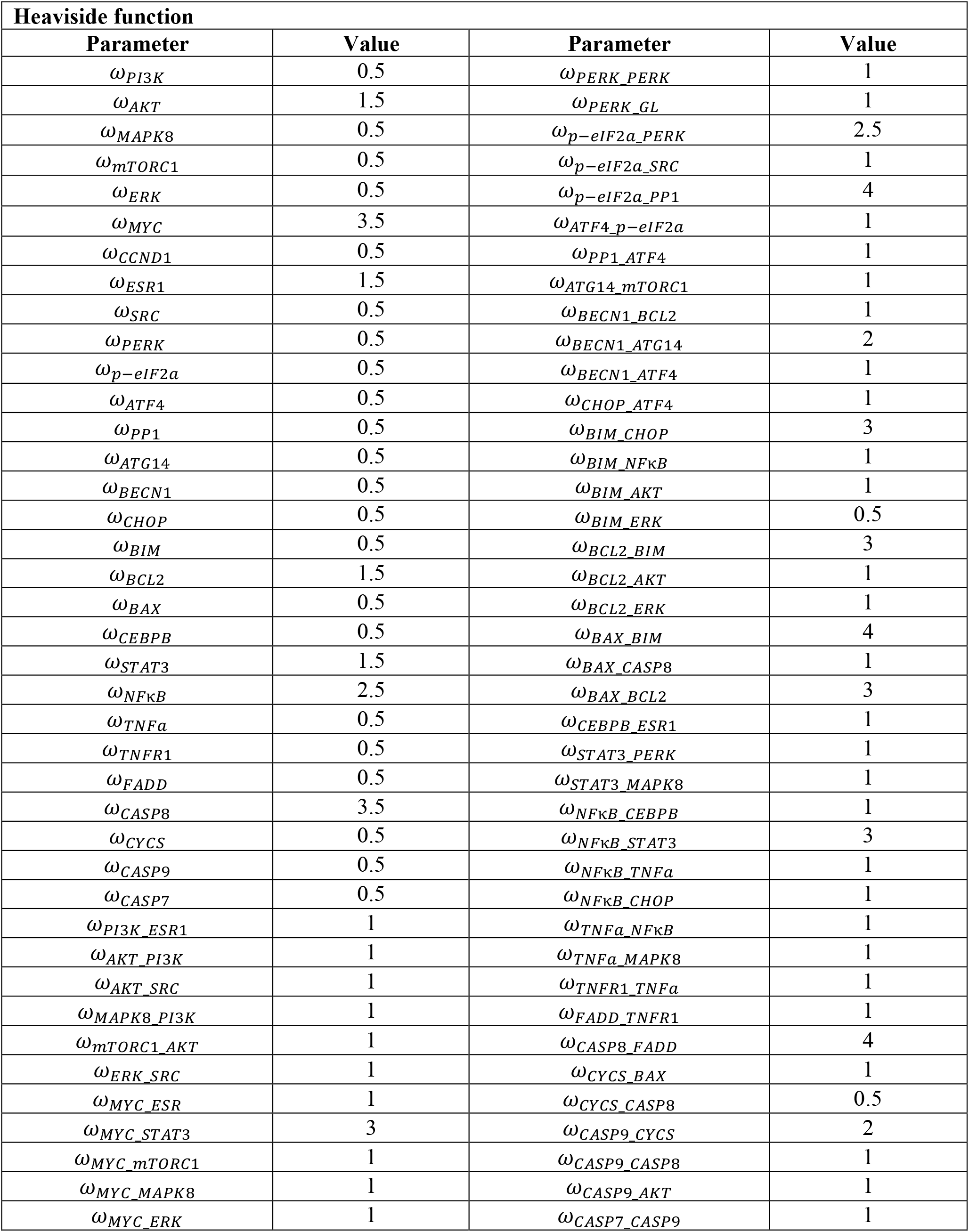
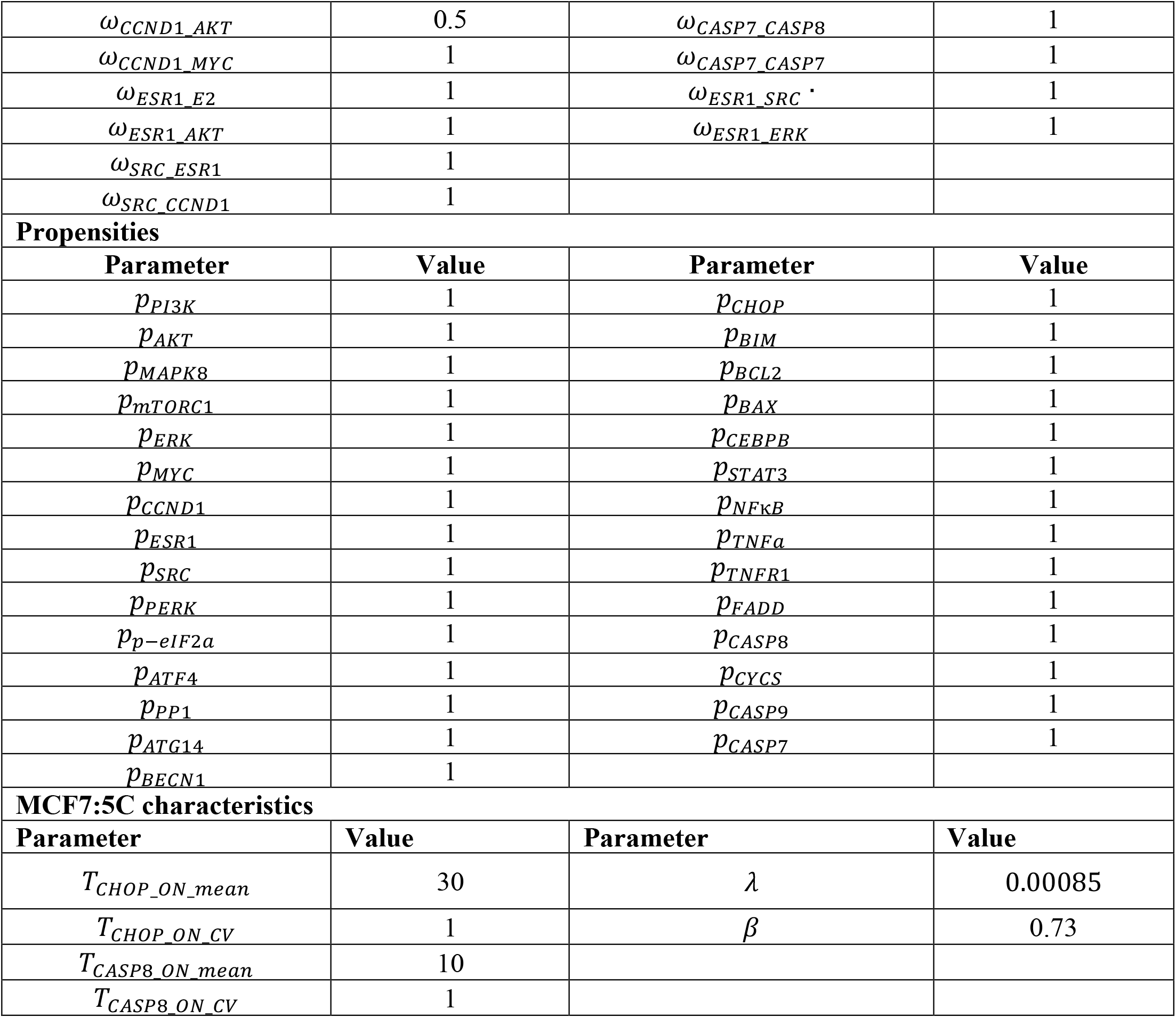
The model parameter values.

## References

1. Feng, Y. et al. Breast cancer development and progression: risk factors, cancer stem cells, signaling pathways, genomics, and molecular pathogenesis. Genes Dis. 5, 77–106 (2018).

2. Rozeboom, B., Dey, N. & De, P. ER+ metastatic breast cancer: past, present, and a prescription for an apoptosis-targeted future. Am. J. Cancer Res. 9, 2821–2831 (2019).

3. Osborne, C. K. Tamoxifen in the treatment of breast cancer. N. Engl. J. Med. 339, 1609–18 (1998).

4. Patel, H. K. & Bihani, T. Selective estrogen receptor modulators (SERMs) and selective estrogen receptor degraders (SERDs) in cancer treatment. Pharmacol. Ther. 186, 1–24 (2018).

5. Sood, A., Lang, D. K., Kaur, R., Saini, B. & Arora, S. Relevance of aromatase inhibitors in breast cancer treatment. Curr. Top. Med. Chem. 21, 1319–1336 (2021).

6. Cuzick, J. et al. Use of anastrozole for breast cancer prevention (IBIS-II): long-term results of a randomised controlled trial. Lancet 395, 117–122 (2020).

7. Fisher, B. et al. A randomized clinical trial evaluating tamoxifen in the treatment of patients with node-negative breast cancer who have estrogen-receptor-positive tumors. N. Engl. J. Med. 320, 479–84 (1989).

8. Vasan, N., Baselga, J. & Hyman, D. M. A view on drug resistance in cancer. Nature 575, 299–309 (2019).

9. Fisusi, F. A. & Akala, E. O. Drug combinations in breast cancer therapy. Pharm. Nanotechnol. 7, 3–23 (2019).

10. Fisher, B. et al. Tamoxifen and chemotherapy for lymph node-negative, estrogen receptor-positive breast cancer. J. Natl. Cancer Inst. 89, 1673–82 (1997).

11. Hughes, J. P., Rees, S., Kalindjian, S. B. & Philpott, K. L. Principles of early drug discovery. Br. J. Pharmacol. 162, 1239–49 (2011).

12. Li, H., Xuan, J., Wang, Y. & Zhan, M. Inferring regulatory networks. Front. Biosci. 13, 263–75 (2008).

13. Gómez Tejeda Zañudo, J. et al. Cell line-specific network models of ER+ breast cancer identify potential PI3Kα inhibitor resistance mechanisms and drug combinations. Cancer Res. 81, 4603–4617 (2021).

14. Gómez Tejeda Zañudo, J., Scaltriti, M. & Albert, R. A network modeling approach to elucidate drug resistance mechanisms and predict combinatorial drug treatments in breast cancer. Cancer Converg. 1, 5 (2017).

15. Taoma, K., Ruengjitchatchawalya, M., Liangruksa, M. & Laomettachit, T. Boolean modeling of breast cancer signaling pathways uncovers mechanisms of drug synergy. PLoS One 19, e0298788 (2024).

16. Tsirvouli, E. et al. A middle-out modeling strategy to extend a colon cancer logical model improves drug synergy predictions in epithelial-derived cancer cell lines. Front. Mol. Biosci. 7, 502573 (2020).

17. Vitali, F. et al. A network-based data integration approach to support drug repurposing and multi-target therapies in triple negative breast cancer. PLoS One 11, e0162407 (2016).

18. Ellis, M. J. et al. Lower-dose vs high-dose oral estradiol therapy of hormone receptor-positive, aromatase inhibitor-resistant advanced breast cancer: a phase 2 randomized study. JAMA 302, 774–80 (2009).

19. Lewis-Wambi, J. S. & Jordan, V. C. Estrogen regulation of apoptosis: how can one hormone stimulate and inhibit? Breast Cancer Research 11, 1–12 (2009).

20. Lewis, J. S., Osipo, C., Meeke, K. & Jordan, V. C. Estrogen-induced apoptosis in a breast cancer model resistant to long-term estrogen withdrawal. J. Steroid Biochem. Mol. Biol. 94, 131–41 (2005).

21. Sengupta, S., Sevigny, C. M., Bhattacharya, P., Jordan, V. C. & Clarke, R. Estrogen-induced apoptosis in breast cancers is phenocopied by blocking dephosphorylation of eukaryotic initiation factor 2 alpha (eIF2α) protein. Mol. Cancer Res. 17, 918–928 (2019).

22. Fan, P. et al. Modulation of nuclear factor-kappa B activation by the endoplasmic reticulum stress sensor PERK to mediate estrogen-induced apoptosis in breast cancer cells. Cell Death Discov. 4, 15 (2018).

23. Fan, P. et al. Integration of downstream signals of insulin-like growth factor-1 receptor by endoplasmic reticulum stress for estrogen-induced growth or apoptosis in breast cancer cells. Mol. Cancer Res. 13, 1367–76 (2015).

24. Dagogo-Jack, I. & Shaw, A. T. Tumour heterogeneity and resistance to cancer therapies. Nat. Rev. Clin. Oncol. 15, 81–94 (2018).

25. Harbeck, N. et al. Breast cancer. Nat. Rev. Dis. Primers 5, 66 (2019).

26. Lewis, J. S. et al. Intrinsic mechanism of estradiol-induced apoptosis in breast cancer cells resistant to estrogen deprivation. J. Natl. Cancer Inst. 97, 1746–59 (2005).

27. Jordan, V. C. The new biology of estrogen-induced apoptosis applied to treat and prevent breast cancer. Endocr. Relat. Cancer 22, R1–R31 (2015).

28. Lo Surdo, P. et al. SIGNOR 4.0: the 2025 update with focus on phosphorylation data. Nucleic Acids Res. 54, D682–D690 (2026).

29. Laomettachit, T., Kraikivski, P. & Tyson, J. J. A continuous-time stochastic Boolean model provides a quantitative description of the budding yeast cell cycle. Sci. Rep. 12, 20302 (2022).

30. Fan, P. et al. C-Src modulates estrogen-induced stress and apoptosis in estrogen-deprived breast cancer cells. Cancer Res. 73, 4510–4520 (2013).

31. Obiorah, I., Sengupta, S., Fan, P. & Jordan, V. C. Delayed triggering of oestrogen induced apoptosis that contrasts with rapid paclitaxel-induced breast cancer cell death. Br. J. Cancer 110, 1488–1496 (2014).

32. Lui, A., New, J., Ogony, J., Thomas, S. & Lewis-Wambi, J. Everolimus downregulates estrogen receptor and induces autophagy in aromatase inhibitor-resistant breast cancer cells. BMC Cancer 16, 1–15 (2016).

33. Escher, T. E. et al. Enhanced IFNα signaling promotes ligand-independent activation of ERα to promote aromatase inhibitor resistance in breast cancer. Cancers (Basel). 13, (2021).

34. Soung, Y. H. et al. Inactivating mutations of CASPASE-7 gene in human cancers. Oncogene 22, 8048–52 (2003).

35. Kurokawa, H. et al. Alteration of caspase-3 (CPP32/Yama/apopain) in wild-type MCF-7, breast cancer cells. Oncol. Rep. 6, 33–7 (1999).

36. Bottosso, M. et al. Moving toward precision medicine to predict drug sensitivity in patients with metastatic breast cancer. ESMO Open 9, 102247 (2024).

37. Li, X. et al. Oleandrin, a cardiac glycoside, induces immunogenic cell death via the PERK/elF2α/ATF4/CHOP pathway in breast cancer. Cell Death Dis. 12, 314 (2021).

38. Ge, A. et al. Mechanism of ferroptosis in breast cancer and research progress of natural compounds regulating ferroptosis. J. Cell. Mol. Med. 28, e18044 (2024).

39. Chen, C. et al. Targeting pyroptosis in breast cancer: biological functions and therapeutic potentials on It. Cell Death Discov. 9, 75 (2023).

40. Yan, J., Wan, P., Choksi, S. & Liu, Z.-G. Necroptosis and tumor progression. Trends Cancer 8, 21–27 (2022).

